# Batch effects removal for microbiome data via conditional quantile regression (ConQuR)

**DOI:** 10.1101/2021.09.23.461592

**Authors:** Wodan Ling, Ni Zhao, Anju Lulla, Anna M. Plantinga, Weijia Fu, Angela Zhang, Hongjiao Liu, Zhigang Li, Jun Chen, Timothy Randolph, Wei Li A. Koay, James R. White, Lenore J. Launer, Anthony A. Fodor, Katie A. Meyer, Michael C. Wu

## Abstract

Batch effects in microbiome data arise from differential processing of specimens and can lead to spurious findings and obscure true signals. Most existing strategies for mitigating batch effects rely on approaches designed for genomic analysis, failing to address the zero-inflated and over-dispersed microbiome data. Strategies tailored for microbiome data are restricted to association testing, failing to allow other analytic goals such as visualization. We develop the Conditional Quantile Regression (ConQuR) approach to remove microbiome batch effects using a two-part quantile regression model. It is a fundamental advancement in the field because it is the first comprehensive method that accommodates the complex distributions of microbial read counts, and it generates batch-removed zero-inflated read counts that can be used in and benefit all usual subsequent analyses. We apply ConQuR to real microbiome data sets and demonstrate its state-of-the-art performance in removing batch effects while preserving or even amplifying the signals of interest.

---

Advances in 16S rRNA^1^ and full genome^2^ sequencing technologies have enabled large-scale human microbiome profiling studies involving hundreds to thousands of individuals. The large sample sizes of these studies and the rich availability of metadata promise a comprehensive understanding of the role of microorganisms in health and disease. These studies have already revealed associations between bacterial taxa and both outcomes and exposures, such as obesity^3^, type 2 diabetes^4^, bacterial vaginosis^5^, antibiotics^6^, and environmental pollutants^7^. However, although the large-scale studies facilitate more robust and powerful analyses, they are often subject to serious batch effects – systematic variation in the data originating from differential handling and processing of specimens^8^. Many large studies include samples collected across times or locations and processed in different runs. In a more extreme situation, several studies may be pooled together for integrative analysis, with inter-study heterogeneity introducing even more severe variation. These batch effects pose a grand challenge and can simultaneously lead to excessive false positive discoveries, obscure true associations between microbes and clinical variables, and hinder prediction modeling and biomarker development. Unfortunately, despite the importance of batch effects, relatively little has been done on mitigating batch effects for microbiome data.

Batch effects are not unique to microbiome data^9^, and standard tools have been developed for other genomic technologies, with the most commonly applied approach being ComBat^10^. However, ComBat and related methods that remove genomic batch effects assume continuous, normally distributed outcomes. Extensions for count data^11^ exist but still make restrictive distributional assumptions. Microbiome data are usually highly zero-inflated, over-dispersed, and heterogeneous with complex distributions. Thus, methods from the other contexts cannot adequately address these issues. At the same time, the limited work on batch effects correction tailored for microbiome data^12–14^ can only be used for batch adjustment in association testing, sometimes even requiring specific types of controls/spike-ins. These approaches fail to allow other designs and other common analytic goals such as visualization. We emphasize that in this paper, batch removal refers to disentangling the batch effects that could otherwise contaminate the signal of key variables and generate batch-removed data that are suitable for any subsequent analyses, while batch adjustment means including batch ID as a covariate in testing. Recently, MMUPHin^15^ extended ComBat to microbiome analysis by considering zero inflation. But ultimately, it assumes the data to be zero-inflated Gaussian, which is only appropriate for certain microbiome data transformations, making many other analyses (e.g., visualization using UniFrac distance) challenging. Therefore, more flexible approaches are direly needed.

In this paper, we propose the conditional quantile regression (ConQuR) approach, the first comprehensive microbiome batch effects removal tool. It works directly on taxonomic read counts and generates corrected read counts that enable all usual microbiome analyses (visualization, association analysis, prediction, etc.) with few restrictions. ConQuR assumes that for each microorganism, samples share the same conditional distribution if they have identical intrinsic characteristics (with the same values of key variables and important covariates, e.g., clinical, demographic, genetic, and other features), regardless of in which batch they were processed. This does not mean the samples have identical observed values, but they share the same distribution for that microbe. Then operationally, for each taxon and each sample, ConQuR non-parametrically models the underlying distribution of the observed value, adjusting for key variables and covariates, and remove the batch effects relative to a chosen reference batch. It is fundamentally different from quantile normalization, the widely used approach to align gene expression data. ConQuR allows various distributions for different taxa and works on the conditional distributions depending on metadata, while quantile normalization assumes all taxa are homogeneous and makes the empirical marginal distribution identical to a reference batch. Moreover, ConQuR is fundamentally different from the genomic batch removal methods. Instead of using parametric models, ConQuR uses a composite non-parametric model to correct the entire complex conditional distribution of microbial read counts, robustly and thoroughly, while maintaining the zero-inflated integer nature of microbiome data. In particular, we use quantile regression for counts^16, 17^ to model the read counts among samples for which the microbe is present, and separately model the presence-absence status of the microbe by logistic regression. Quantile regression is non-parametric and directly models percentiles of the outcome, such as the median and quartiles. ConQuR is, therefore, robust to microbiome data characteristics and able to correct higher-order batch effects beyond the mean and variance differences. With zeros explicitly modeled by logistic regression, ConQuR can also address the batch variation affecting the presence-absence status of microbes.

To systematically evaluate ConQuR, we study two large microbiome data sets with different types of batch effects. One data set comes from a single large gut microbiome study of cardiovascular diseases and has moderate batch differences with samples sequenced across several runs. The second one suffers from more substantial “batch” effects as the data were collected and sequenced in different HIV gut microbiome studies. By visual and numerical comparisons, we demonstrate that ConQuR thoroughly removes the batch effects, and preserves the effects of key variables in both association testing and prediction. All usual data transformations and analyses can be conducted on the corrected read count data with minimal regard for the batches.

## Results

### Overview of ConQuR

The central objective of ConQuR is to remove batch effects while preserving real signals. This is done on a taxon-by-taxon and sample-by-sample basis using a two-step procedure (Fig. 1a). In the first regression-step, we regress out the batch effects using a non-parametric extension of the two-part model^18^ for zero-inflated count outcomes. Specifically, a logistic model determines the likelihood of the taxon’s presence, and quantile regression models percentiles of the read count distribution given the taxon is present. The explanatory variables include the batch ID, key variables, and important covariates based on prior knowledge. Accordingly, we can robustly estimate the entire original distribution of the taxon count for each sample, and also estimate the batch-free distribution by subtracting the fitted batch effects relative to a chosen reference batch from both the logistic and quantile parts. Note that we fit the two-part model using all samples for a particular taxon, but due to differences in sample characteristics, the conditional distributions are sample specific. In the second matching-step (Fig. 1b), we locate the sample’s observed count in the estimated original distribution, and then pick the value at the same percentile in the estimated batch-free distribution as the corrected measurement. We repeat this two-step correction for each sample and then each taxon. More details are described in the “Methods” section.

**Fig. 1.**
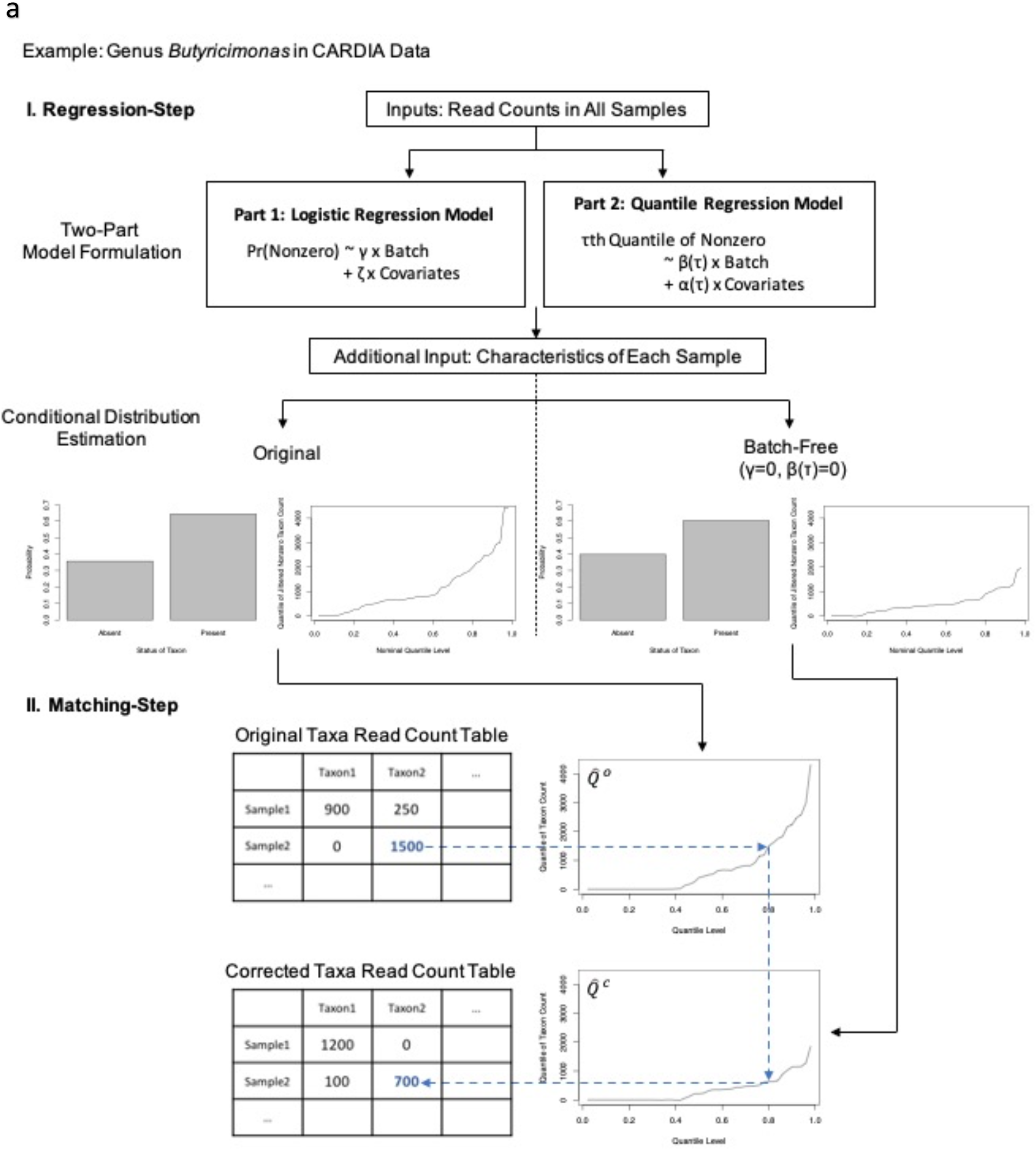
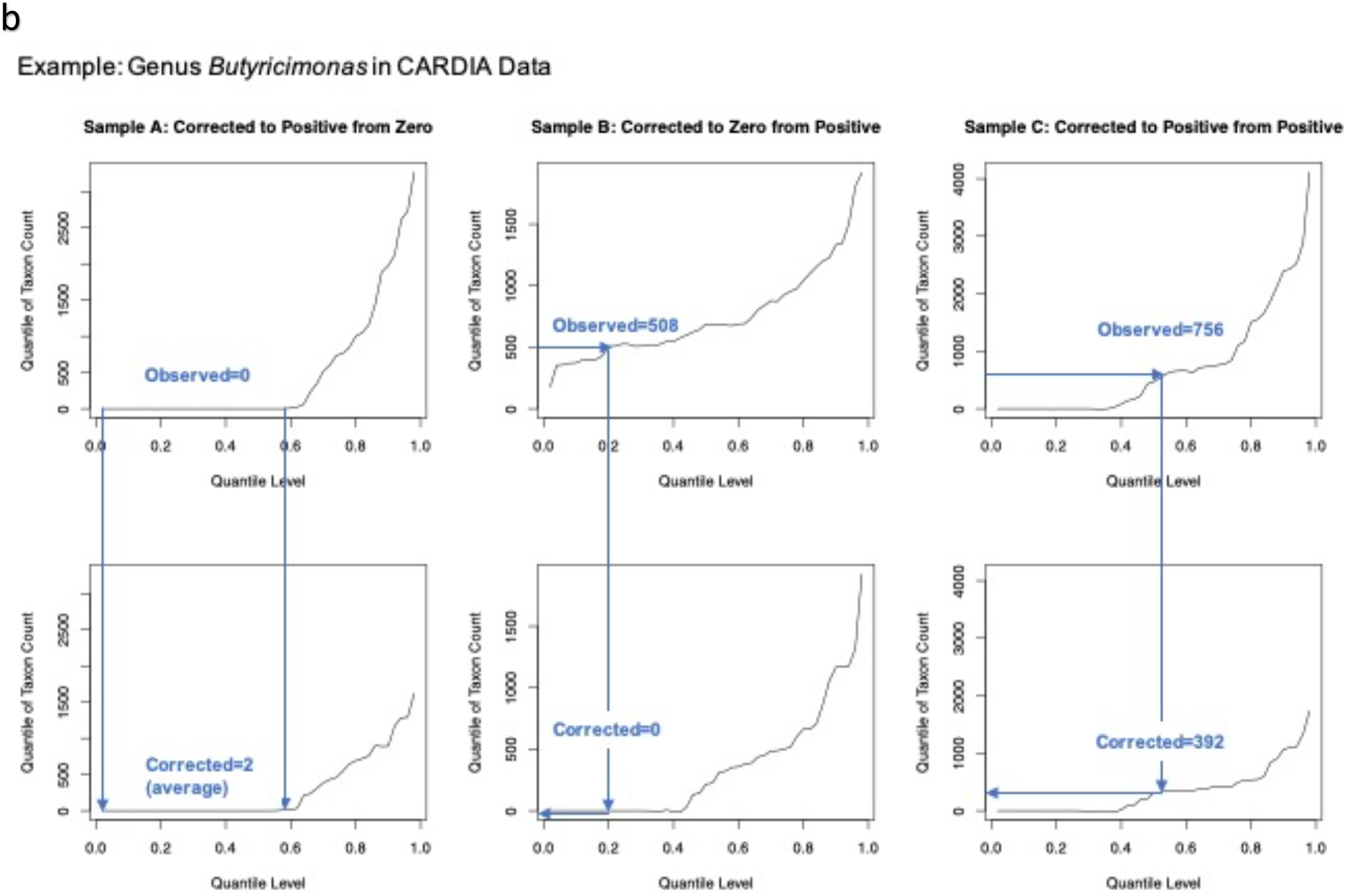
Illustration of the ConQuR algorithm. **a**, Two-step procedure: I. regression-step: (1) Use all available samples to fit the two-part quantile regression model; (2) For each sample, estimate the original likelihood of the taxon being present and the original distribution (by estimating a fine grid of percentiles) given the taxon is present. The two parts jointly determines the zero-inflated, over-dispersed conditional quantile function (the inverse of conditional distribution function) of the taxon count 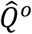. In the same manner, estimate the batch-free conditional quantile distribution 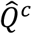. II. Matching-step: locate the observed read count in 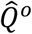, and pick the value at the same location of 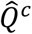 as the corrected read count. Repeat the procedure for each sample and then each taxon. **b**, Three scenarios of matching: (left panel) Sample A has a less sparse and less outlying estimated batch-free distribution compared to the original one, its observed measurement zero is corrected to be a non-zero number, (middle panel) Sample B has a sparser and more outlying estimated batch-free distribution than the original one, its observed non-zero count, located at a lower percentile of the original distribution, is corrected to be zero, (right panel) Sample C has a slightly less sparse and less outlying estimated batch-free distribution than the original one, its observed non-zero count, located at a middle percentile of the original distribution, is corrected to be a smaller non-zero count.

The modeling and estimation framework of ConQuR has four advantages. First, as it directly estimates every conditional percentile without specific assumptions, the complex microbial count distribution is robustly and comprehensively captured. It is more reliable (robust and flexible) than a parametric model, such as negative binomial or Gaussian, which requires the read count to follow a specific shape. Second, the composite model of logistic and quantile regressions allows heterogeneous associations between the zero-inflated over-dispersed microbial counts and traits, i.e., batch effects do not need to be uniform across the range of the taxon’s abundance. Consequently, the batch effects removal is thorough, mitigating the mean, variance, and even higher-order batch effects. Finally, as the framework handles zero inflation, it calibrates the unwanted presence-absence differences among batches, thereby recovering non-zero numbers for under-sampled observations and forcing those over-sampled to be zeros.

### Application to a single large-scale epidemiology study

We examine whether ConQuR can mitigate the batch effects via application to real data. We first applied it to a study containing traditional batch variation: samples are collected under one protocol but handled in different batches. The Coronary Artery Risk Development in Young Adults (CARDIA) Study^19^ enrolled young adults in 1985-86, with the aim of elucidating the development of cardiovascular disease (CVD) risk factors across adulthood. A variety of factors related to CVD have been collected, including health behaviors, medical history (e.g., medication use), and clinical risk factors, such as blood pressure. Basic demographic measures such as age, gender, and race have also been collected. At the Year 30 follow-up examination (2015-2016), stool samples were collected and processed for DNA extraction and library preparation across four batches. Then, the 16S rRNA marker gene (V3-V4) was sequenced by Illumina technology (MiSeq 2×300) over seven runs (approximately 96 samples/run), two from each of the first three DNA extraction batches, and the last run from the fourth batch. Thus, at the finest level, data were generated across seven batches. Following sequencing, forward reads were processed through the DADA2^20^ pipeline for quality control and derivation of amplicon sequence variants (ASVs), and taxonomy was assigned using the Silva reference database^21^. The data were aggregated to the genus level, and lineages with zero reads across all samples were excluded.

Batch ID (Batches 0 to 6) indicates in which of the seven sequencing runs each sample was included. Systolic blood pressure (SBP) was the primary variable of interest. Covariates considered for adjustment included gender (Female = 1, Male = 0) and race (Black = 1, White = 0). With missingness filtered out, the final-processed data included 375 genera from 633 samples (Supp. Tab. 1). We aimed to remove the effects of other batches relative to Batch 3, assuming that SBP, gender, and race could jointly describe the conditional distribution for each sample of each taxon’s abundance. Although there are no methods tailored for microbiome batch effects removal on the read count scale, we included ComBat-seq^11^, which was developed for RNA-seq data, as a potential comparator.

We first demonstrated the efficacy of ConQuR through visualization: PCoA plots with colors representing batch IDs. We used Bray-Curtis dissimilarity on count data, Euclidean dissimilarity on the corresponding centered log-ratio (CLR)^22, 23^ transformed relative abundance data (Aitchison dissimilarity), and GUniFrac dissimilarity (a compromise between unweighted UniFrac distance and weighted UniFrac distance, which is computed based on relative abundance). As Fig. 2a shows, in terms of all three dissimilarities, the un-corrected data exhibited significant differences among batches, and ConQuR performed a thorough correction in both the mean (centroids) and dispersion (sizes of ellipses). Specifically, in the raw count scale (by Bray-Curtis distance), ConQuR centered the means of the seven batches to the same point. As an ellipse connects the 95% percentile of points for each batch in the bivariate plot, we see that ConQuR not only made the extent of variability almost the same across batches but also removed higher-order effects of batch (angles of the ellipses). In the relative abundance scale (by Aitchison or GUniFrac distances), ConQuR also succeeded in aligning the different batches. However, its advantage over ComBat-seq was not as substantial as in the raw count scale. This is because ComBat-seq includes the library size as an offset. It, in fact, models the mean and dispersion of relative abundance. To examine ConQuR’s performance on taxa with different frequencies, we created PCoA plots on common and rare genera separately. We see that ConQuR showed dominating performance on normal to common taxa that are present in more than 50% of the samples and demonstrated comparable correction on the rare ones compared to ComBat-seq (Supp. Fig. 3).

**Fig. 2.**
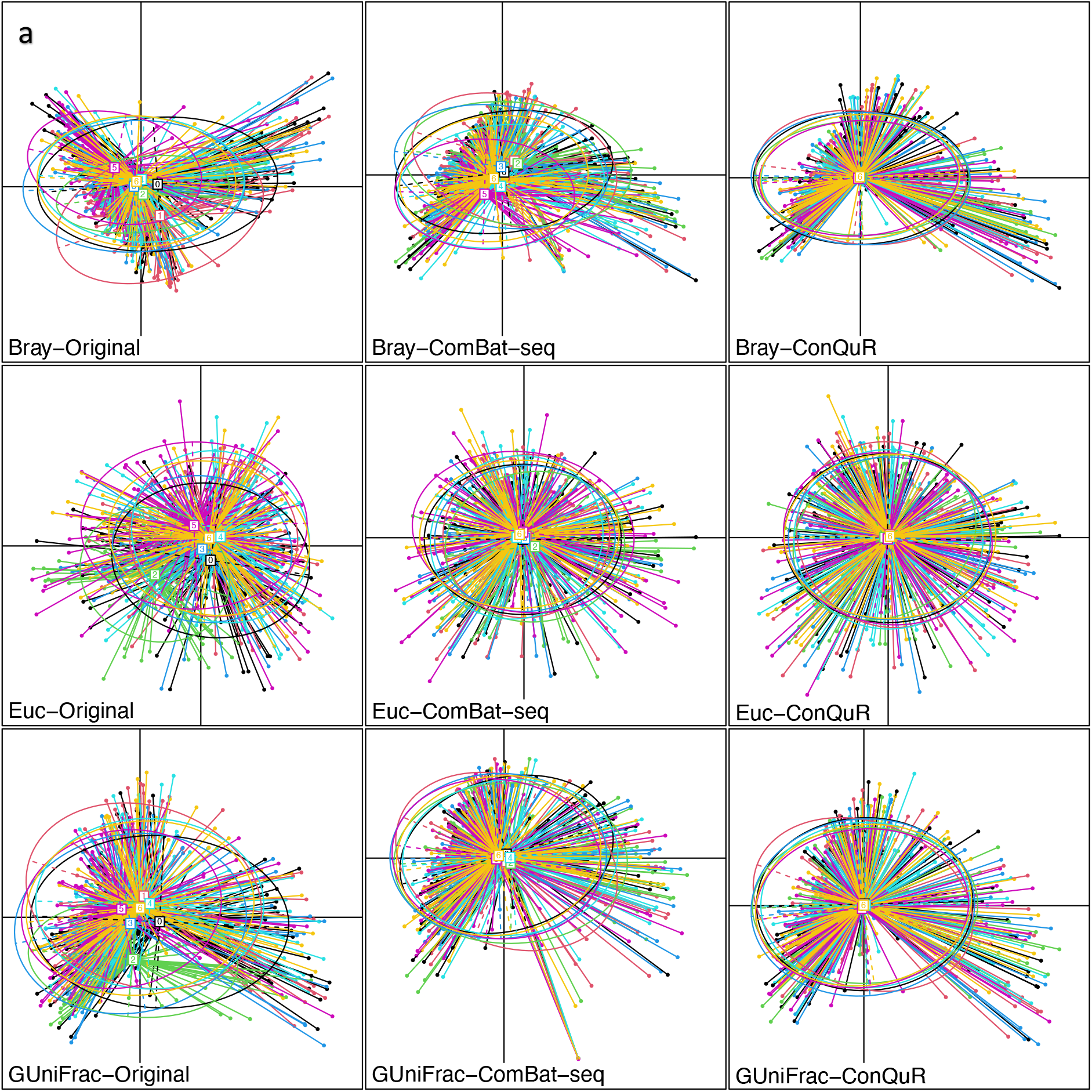
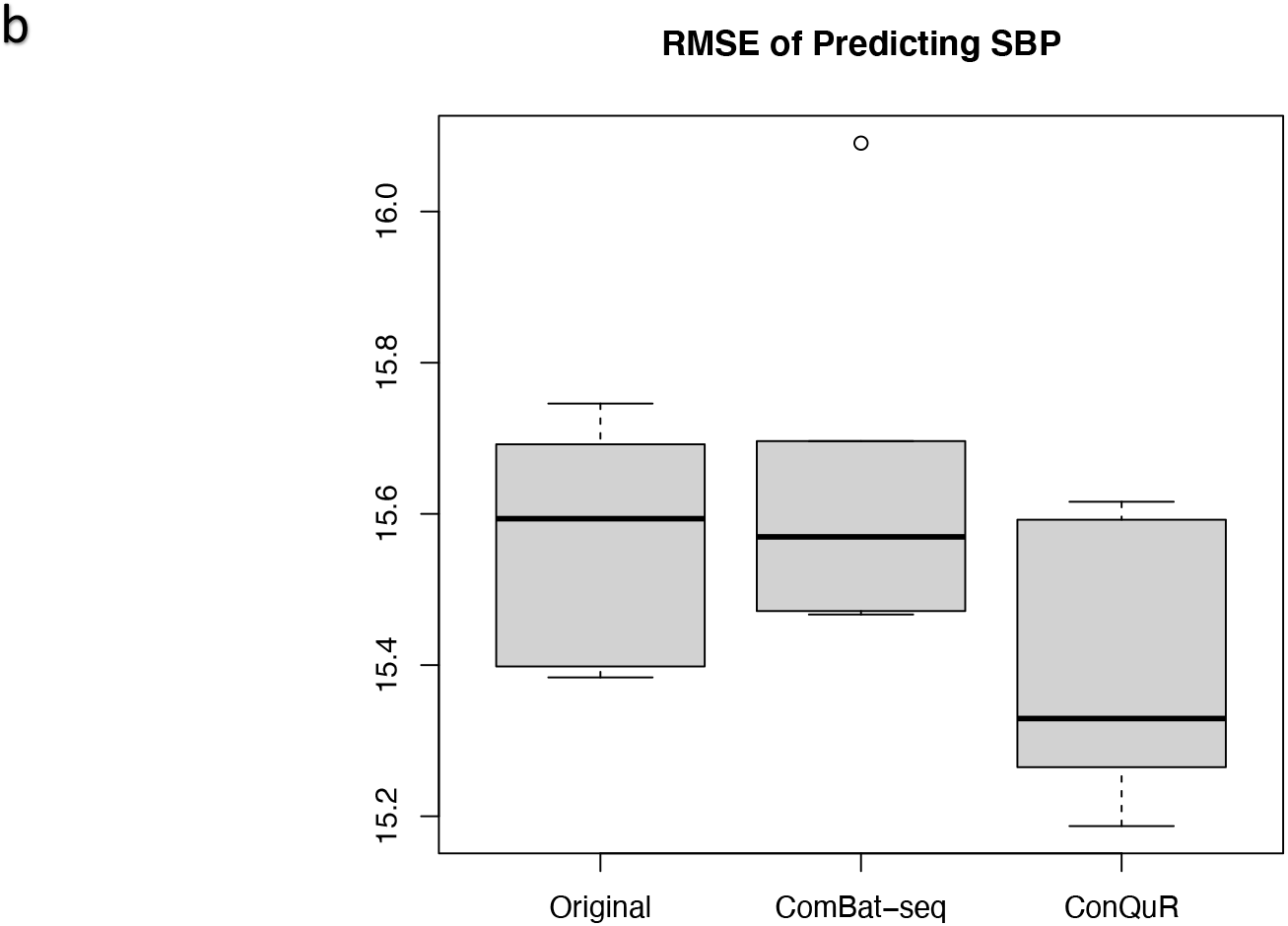
Visualization and prediction metric evaluated on the original and corrected CARDIA data. **a**, PCoA plots clustered by batch ID,(top panel) by Bray-Curtis distance on raw count scale of the original data, ComBat-seq corrected data, and ConQuR corrected data, (middle panel) by Euclidean distance on the corresponding CLR transformed relative abundance scale of the data, and (bottom panel) by GUniFrac distance. **b**, RMSE of predicting systolic blood pressure (SBP), based on (left) the original data, (middle) ComBat-seq corrected data, and (right) ConQuR corrected data, via random forest models across five folds.

Further, we numerically evaluated ConQuR’s ability to remove batch effects while preserving the signals of interest. First, we assessed the variability of the microbiome data explained by batch ID and the key variable SBP, using PERMANOVA^24^ *R*^2^ (in Bray-Curtis dissimilarity). Note that as a measure of multivariate correlation, there is no easy interpretation of PERMANOVA *R*^2^; nonetheless, it is a reliable metric to quantify the proportion of data variability explained by a particular variable. As Tab. 1 shows, batch explained 5.66% of the variability in the original CARDIA count data. ConQuR reduced the proportion to 0.10%. At the same time, it maintained the amount of variability explained by SBP (0.38% vs. 0.37% in the original data). After normalizing the corrected read counts into relative abundance scale, we still observed the improvement by ConQuR – reducing the variability explained by batch from 3.65% to 0.25% and keeping the explanatory power of SBP (0.36% vs. 0.34% in the original data). Given the modest PERMANOVA *R*^2^ values, we augmented the numerical results by examining the prediction ability of the corrected taxa read count table. Random forest was chosen for the task to allow for flexible and non-linear modeling. Five-fold cross-validation on the root of mean squared error (RMSE) of predicting SBP was used to evaluate the accuracy. As this analysis merely investigates the performance of ConQuR via an alternative metric, we applied ConQuR to the combined training and testing sets for simplicity. Also, being a completely different metric, how accurate the taxa read count table predicts SBP is not inferable by the multivariate correlation PERMANOVA *R*^2^. We used boxplots to summarize the results across folds (Fig. 2b). ConQuR systematically lowered the RMSE, amplifying the accuracy of predicting SBP from the microbial profiles. Collectively, numerical evaluations demonstrate that ConQuR maintains the signals of interest while thoroughly removing batch effects, enabling more reliable community-level association testing (by PERMANOVA or MiRKAT^25^, a generalization of PERMANOVA) and more accurate prediction.

**Tab. 1.** PERMANOVA *R*^2^ (in Bray-Curtis dissimilarity) explained by batch ID and systolic blood pressure (SBP) in the original and corrected CARDIA data. (left) in the raw count scale and (right) in the corresponding relative abundance scale.

|  | Raw count |  | Relative abundance |  |
| --- | --- | --- | --- | --- |
|  | batch | SBP | batch | SBP |
| Original | 0.0566 | 0.0037 | 0.0365 | 0.0034 |
| ComBat-seq | 0.0260 | 0.0032 | 0.0070 | 0.0030 |
| ConQuR | 0.0010 | 0.0038 | 0.0025 | 0.0036 |

A recurring objective in microbiome studies is the implication of individual taxa; thus, we finally examined the association between individual taxa and SBP before and after ConQuR correction. To make the analysis conservative and general, we used ordinary linear regression of taxon relative abundance on SBP, adjusting for gender and race. The false discovery rate (FDR) was controlled by Benjamini–Hochberg (BH) procedure at *α*=0.05. No genus was found to be significantly associated with SBP in the original or ComBat-seq corrected data. In contrast, *Anaerovoracaceae_Family_XIII_UCG-001* (adjusted p-value=0.0012) and *Hydrogenoanaerobacterium* (adjusted p-value=0.0422) were detected to be differentially abundant in the ConQuR corrected data. For adolescents, change in *Family_XIII_UCG-001*’s relative abundance is positively related to changes in triglycerides, serum cholesterol, and low-density lipoprotein cholesterol^26^, which are factors closely associated with hypertension^27, 28^. Also, it is differentially abundant between control and coronary artery disease (CAD) patients^29^, where the strong link between hypertension and CAD has been shown^30, 31^. *Hydrogenoanaerobacterium* is a crucial contributor to modeling the change of blood pressure in studying the effect of fasting on high blood pressure in metabolic syndrome patients^32^. Supported by the biological findings, we confirm that ConQuR helps to peel off the confounding batch effects, maintain the true signals and lead to meaningful discoveries.

### Application to integration of multiple individual studies

We further consider the performance of ConQuR within the context of vertical data integration wherein interest is in combining multiple individual studies. We applied it to data from the HIV re-analysis consortium (HIVRC)^33^. Raw 16s rRNA gene sequencing data from distinct studies were processed through a common pipeline – Resphera Insight^34^. Details of data pre-processing and taxonomic assignment are in^33^. In this paper, we focused on the data aggregated to the genus level. HIV status (Positive = 1, Negative = 0) was regarded as the primary metadata, while age, gender (Female = 1, Male = 0) and BMI were considered as covariates. Retaining complete cases only, we obtained the final data that consists of 606 genera on 572 individuals from 10 studies (Supp. Tab. 2) and regarded Study 0 as the reference batch.

Here, the batch effects are represented by the studies and much more extreme since the studies varied in terms of experimental designs and sequencing protocols (Supp. Tab. 2 of^33^). Measured by PERMANOVA *R*^2^, the study ID explains 30.39% of the data variability, while the traditional batch effects in CARDIA only contribute 5.66%. We also observed substantial imbalance, sparsity, and heterogeneity in the microbial profiles, as they are unlikely to be fully matched across studies. Comparing Supp. Tab. 2 to Supp. Tab. 1, we see that only 65 out of the 606 genera are present in all studies, while the ratio is 183/375 in CARDIA. The library size ranges quite differently across studies, e.g., samples have 185-1000 reads in Study 6, whereas the library size was rarefied to 20000 reads in Studies 0 and 8. Note that we intentionally kept the samples with minimal library sizes to show ConQuR’s capability to handle the outliers. Correcting such data would be more challenging than a relatively homogeneous one like the CARDIA data. The imbalance in metadata (sample sizes and characteristics, Supp. Tab. 2) also adds to the difficulty of batch effects removal.

Again, we first used PCoA plots grouped by the study ID to examine ConQuR visually. Since we did not obtain a phylogenetic tree for the pooled HIVRC data, we omitted the plots based on GUniFrac dissimilarity. From Fig. 3a, we see that ConQuR considerably removed the study variation in the raw count data (by Bray-Curtis distance). The means of the 10 studies (centroids) came almost together. Their dispersions and higher-order features (sizes and angles of the ellipses) are much more aligned with each other compared to the original data. On the corresponding relative abundance data, though ConQuR did not demonstrate perfect correction, it still made the 10 studies substantially more harmonized – brought the means far closer than the original data, also amplified the dispersions of the minimally variable studies, e.g., Study 6, making its variance approximate to the others. Again, ConQuR showed a more thorough correction on genera with more than 50% prevalence, and was non-inferior on rare genera, as compared to ComBat-seq (Supp. Fig. 4).

**Fig. 3.**
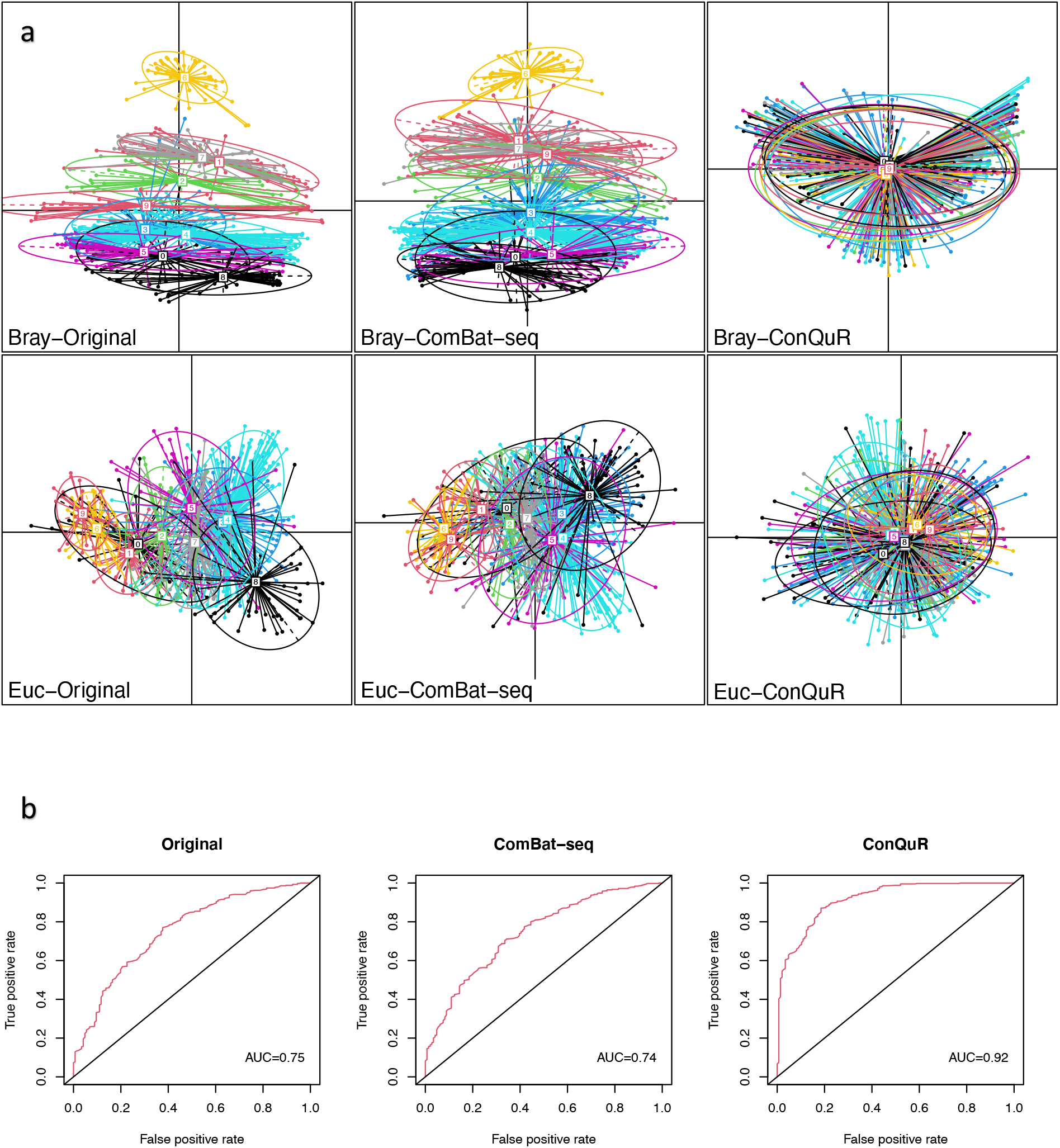
Visualization and prediction metric evaluated on the original and corrected HIVRC data. **a**, PCoA plots clustered by study ID, (top panel) by Bray-Curtis distance on raw count scale of the original data, ComBat-seq corrected data, and ConQuR corrected data, (bottom panel) by Euclidean distance on the corresponding CLR transformed relative abundance scale of the data. **b**, ROC curve and AUC of predicting HIV status, based on (left) the original data, (middle) ComBat-seq corrected data, and (right) ConQuR corrected data, via random forest models across five folds.

Next, we used PERMANOVA *R*^2^ to examine ConQuR numerically. Although ConQuR did not make the correction as perfect as on the traditional batch sequencing microbiome data, it maintained its effectiveness in terms of the proportion of unwanted variation eliminated. For the CARDIA data, ConQuR reduced 98% batch effects from 5.66% to 0.10. While for the HIVRC data in the raw count scale, ConQuR again mitigated 94% of the variation due to study from 30.39% to 1.94% (Tab. 2). Meanwhile, ConQuR preserved the variability explained by HIV status (0.59% vs. 0.57% in the original data). For the normalized data in the relative abundance scale, ConQuR decreased the percentage of study-explained variability from 18.66% to 1.63%. Then, we predicted the binary HIV status using random forests, based on the original and corrected HIVRC data. We computed the average area under (AUC) the receiver operating characteristic (ROC) curve across five folds, concatenated the prediction results, and plotted the overall ROC curve. As Fig. 3b shows, ConQuR boosted the average AUC from 0.75 to 0.92. Overall, the numerical evaluations show that ConQuR is robust to different types of batch effects, demonstrating stable performance in thoroughly mitigating batch variation while keeping signals of interest, even when the batches are highly heterogeneous.

**Tab. 2.** PERMANOVA *R*^2^ (in Bray-Curtis dissimilarity) explained by study ID and HIV status in the original and corrected HIVRC data. (left) in the raw count scale and (right) in the corresponding relative abundance scale.

|  | Raw count |  | Relative abundance |  |
| --- | --- | --- | --- | --- |
|  | study | HIV status | study | HIV status |
| Original | 0.3039 | 0.0057 | 0.1866 | 0.0085 |
| ComBat-seq | 0.2064 | 0.0063 | 0.0710 | 0.0092 |
| ConQuR | 0.0194 | 0.0059 | 0.0163 | 0.0065 |

For the individual-taxon association analysis, linear regression (adjusting for age, gender, and BMI) on the original relative abundance data did not find differentially abundant genera between control and HIV+ patient. On the ComBat-seq corrected data, one genus, *Anaerosporobacter* (adjusted p-value=0.0319), was identified. While on the ConQuR corrected data, three genera were detected: *Acidaminococcus, Parasporobacterium*, and *Vampirovibrio* (adjusted p-values=0.0159, 0.0159, and 0.0161). *Acidaminococcus* has been shown to increase in HIV+ patients^35^. Again, it confirms that ConQuR can disentangle signals from the unwanted variation and increase the power of detecting differentially abundant taxa.

## Discussion

Batch effects removal is a challenging task for microbiome data. Approaches designed for genomic data make strong parametric assumptions, which may be inadequate to address the complex distribution of microbiome data. On the other hand, methods tailored toward microbiome data are restricted to association testing or specialized study designs, failing to provide a corrected read count table for other kinds of subsequent analyses in general studies.

We present ConQuR, a robust approach to thoroughly remove unwanted batch variation in microbiome data and generate corrected taxa read count tables. It is based on a two-part quantile regression framework, simultaneously handling the complex read count distribution by quantile regression and the presence-absence status of microbes by logistic regression. ConQuR is most suited for situations in which the microbiome data are processed from highly heterogenous batches and follow irregular distributions. If interest is only in association analysis, it may be sufficient to use existing microbiome batch adjustment approaches. However, ConQuR represents a powerful option – creating batch-removed taxonomic read counts for more general analyses beyond association, including visualization, prediction, etc.

We applied ConQuR to study two data sets, one from a single cardiovascular study with modest batch effects, and the other from an integrative analysis – sparser, more irregular, and containing more prominent study variation. Visually and numerically, ConQuR demonstrated rigorous performance in correcting the mean, variance, and higher-order batch effects. At the same time, it preserves the effects of key variables in both association and prediction analyses. Finally, ConQuR facilitates relevant biological discoveries about the implication of individual taxa. The principal advantage of ConQuR lies in its non-parametric nature – robust to over-dispersion and heterogeneity, and its capability to handle zero inflation, addressing the complex distributional attributes of microbiome data. All standard down-stream analyses can benefit from it with minimal regard for batches.

Despite the advantages of ConQuR, it has several limitations which are shared by most existing batch removal procedures. First, comprehensive metadata is required to estimate the conditional distributions of read counts accurately. Second, ConQuR uses the metadata twice in both the correction and subsequent analyses, theoretically leading to over-optimism in association analysis^36^. However, in practice, this bias is modest relative to the batch effects, and the inclusion of metadata is often helpful for estimating conditional distributions when the taxon is uncommon or imbalanced among batches. Third, ConQuR cannot work if batch completely confounds the key variable.

In addition to the shared limitations, our results show that ConQuR is imperfect for low frequency taxa. This is because quantile regression cannot provide stable estimates with few non-zero read counts, especially at extremely low or high percentiles. In fact, no methods work very well for those very rare taxa. However, ConQuR still performs better than no correction and improves as sample size increases. Consequently, as microbiome profiling studies continue to increase in size, necessarily inducing more batch effects, the performance of ConQuR will only continue to improve.

## Methods

### Notation

Suppose we have microbiome data from *n* samples, which are sequenced from *B* batches. From each sample, the read counts of *J* taxa are summarized. We therefore have an *n* × *J* table ***Y***, where the entry *Y*_*ij*_ denotes the read count of the *j*th taxon in the *i*th sample. Note that *Y*_*ij*_ is a zero-inflated count variable, and we treat it as the outcome in the proposed regression framework. Besides the batch variable ***B***_*i*_, which is expanded as *B* − 1 dummy variables from the batch ID (the one excluded is the reference batch), each sample has a set of characteristics ***Z***_*i*_. ***Z***_*i*_ includes the key variables of interest in subsequent analyses, important biomedical, demographic, genomic and other information based on prior knowledge, and the intercept. Note that we require the key variables to be included, but do not denote them separately from other covariates when presenting the method, because they play similar roles in the batch effects removal procedure. The concatenated *p*-dimension covariates are denoted by 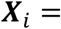 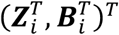. The proposed method is taxon specific, so we henceforth omit the subscript *j* for a simpler presentation.

### Regression-step

One broadly used framework for zero-inflated outcomes is the two-part^18^ or hurdle^37^ model. It separately models the chance that the taxon is present in a sample and the mean of its abundance given it is present. We employ the same strategy, and first assume that the probability of observing a non-zero *Y*_*i*_, *π* = *P*(*Y*_*i*_ > 0|***X***_*i*_), follows a logistic regression model,

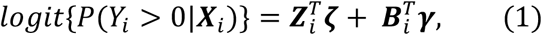

where ***θ***^*L*^ = (***ζ***^*T*^, ***γ***^*T*^)^*T*^ are the true logistic coefficients associated with the covariates and batch variables. Though the presence-absence status of a taxon highly depends on the sequencing depth, we do not explicitly include the depth in the proposed two-part model but automatically deals with it in the logistic part. We assume that batch completely confounds library size (e.g., Batch 1 has mean library size 10000, Batch 2 has mean library size 20000, etc.). Thus, the presence-absence status depends on batch, and we capture the relationship through Model (1). Consequently, one does not need rarefaction before applying ConQuR or include library size as a covariate in the model. As for the variation of library size within a batch, we do not particularly deal with it because (1) usually the variation is not substantial and (2) correcting the between-batch variation is our primary goal.

Next, instead of modeling the mean by traditional parametric models, we use linear quantile regression to model the non-zero part, *Y*_*i*_|*Y*_*i*_ > 0. We assume

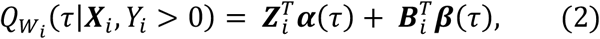

where *W*_*i*_|*Y*_*i*_ > 0 = *Y*_*i*_|*Y*_*i*_ > 0 + *U*, *U* ~ *Uniform*(0,1), and ***θ***^*Q*^(*τ*) = (***α***(*τ*)^*T*^, ***β***(*τ*)^*T*^)^*T*^ are the true quantile coefficients at the *τ*th quantile of *W*_*i*_, which corresponds to a non-zero *Y*_*i*_. Jittering by *U* is a standard technique to apply quantile regression for counts^17^ as it breaks the ties and permits valid estimations and inferences. *τ* is a real value between 0 and 1, indicating the quantile level or percentile, e.g., 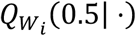 is the conditional median and 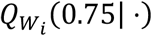 refers to the conditional 3rd quartile of the jittered non-zero read count. Employing a fine grid of quantile levels 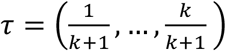 with a large *k* (e.g., 5th, …, 95th percentiles with *k* = 19), we approximately model the conditional quantile function of *W*_*i*_|*Y*_*i*_ > 0. Due to the one-to-one relationship between quantiles of *W*_*i*_|*Y*_*i*_ > 0 and quantiles of *Y*_*i*_|*Y*_*i*_ > 0^17^, the conditional quantile function of *Y*_*i*_ given its presence is established.

We fit the two-part quantile regression model (1)(2) to the investigated taxon with all available samples. Combining the results of the two parts based on the fitted models and a sample’s metadata, we can estimate the original conditional distribution of the taxon count for that sample. As quantile function is the inverse of distribution function, we estimate the conditional quantile function, which is equivalent to estimating the conditional distribution. First, the estimated probability 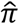 based on the fitted logistic model determines the proportion of zero in the conditional distribution. Specifically, for that sample, all percentiles of the taxon count before the 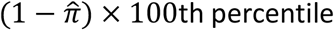 are zero. Next, for percentiles after the 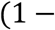 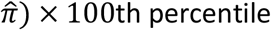, we squeeze in the estimated conditional quantiles of *Y*_*i*_ given its presence. The combined function on the whole probability spectrum (0,1) is the final estimated conditional quantile function 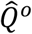, by which the zero-inflated and over-dispersed microbial count distribution is comprehensively revealed. More details of the model and estimation are discussed in the Supplementary Material.

Then, to correct the entire conditional distribution, we regress out batch effects from both the logistic and quantile parts. Specifically, we subtract the estimated effects of batch – ***γ***, and ***β***(*τ*) at any percentile, and then combine the two parts in the same manner to obtain the estimated batch-free conditional quantile function 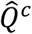. Note that by design, ***γ*** and ***β***(*τ*) are effects of other batches relative to the reference batch (refer to “Notation” section). Therefore, we eliminate the differences between the sample and those in the reference batch having identical characteristics.

### Matching-step

With both the original and batch-free conditional distributions in hand, we find the corresponding value of that sample’s abundance in the batch-free distribution. Ideally, we can find a unique quantile in 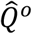 equals to the observed count *Y*_*i*_ (find the 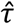 such that 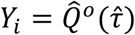). Then, the value at the same percentile in 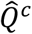 is the corrected read count 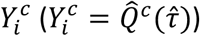.

Since *Y*_*i*_ is a count variable, there might be multiple quantiles in 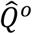 equal to the observed *Y*_*i*_. It is particularly the case when we adjust zero counts, as microbiome data are zero-inflated. By the strict definition of quantiles – *τ* = *P*(*Y*_*i*_ ≤ *y*), we should locate *Y*_*i*_ at the highest percentile where 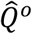 is less or equal to its observed value. For example, in Fig. 1b (left panel), we need to locate the observed zero at the 58th percentile, the rightmost point of the range where 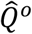 equals zero, and then pick the non-zero count of 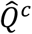 at the same position as the corrected measurement. However, the estimates, particularly around the zero-positive

change point, might not be stable. This is because the estimation of quantile regression is not stable at extreme percentiles. Around the change point, the percentile of non-zero *Y*_*i*_ is approaching the 0th. Therefore, we take the rounded average of all matched quantiles in 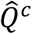 to obtain a “smoothed” 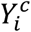, which should be non-zero as well. In this way, we not only allow non-zero values to become zero, but also zero-values to become non-zero.

When the sample size is limited, or the grid of quantile levels is not fine enough, there might be no quantiles in 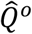 equal to *Y*_*i*_. Then, as quantile functions are left-continuous, we locate *Y*_*i*_ at the percentile with the maximum value smaller than *Y*_*i*_.

After the matching-step, we can correct each sample’s observation for the investigated taxon. Repeat the two-step procedure on all taxa, we adjust all entries in the read count table. As both the presence-absence status and values given the presence of all taxa are corrected relative to the reference batch, we will observe that, in the corrected table, (1) library sizes of other batches are similar to that in the reference batch and (2) non-zero read counts in other batches follow similar distribution as that of the reference.

### Two-layer tuning to achieve the optimal performance

The choice of reference batch affects the quality of 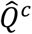, thus also affect the performance of ConQuR. Trying several options is recommended if there is no specific preference based on prior knowledge. Note that using the same reference batch across all taxa is crucial to keep the overall structure of microbiome data. Therefore, tuning over potential reference choices is the top layer of the process.

Instead of the standard logistic and quantile regressions, we can use penalized regressions or drop the key variable and covariates but keep the batch ID only to achieve potentially more stable estimates. Whether the results will benefit from these alternative fitting strategies depends on the specific data set and frequencies of taxa. Therefore, we suggest the second layer of tuning wherein different fitting strategies are used for taxa with different frequencies.

PERMANOVA *R*^2^ explained by batch ID is the tuning criterion, the lower the better. Accordingly, we select the fitting strategy or reference batch that removes batch effects the most. The tuned results are presented in this paper, while visualization of the results by standard ConQuR (using the first batch by numerical or alphabetical order and standard fitting strategy) is included in the Supplementary Material.

#### Tuning over fitting strategies across taxa frequencies

We start with the second layer of tuning, using a pre-specified reference batch.

For the quantile part, one can use L1-penalized quantile regression^38^ with a penalty proportional to sparsity (e.g., 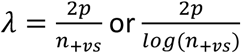). Similar to other LASSO methods, it makes the computation feasible when the non-zero counts are fewer than the number of explanatory variables, and also helps to stabilize the estimates of the conditional quantile functions. A more aggressive alternative is composite quantile regression^39^, which forces different quantiles to share the same set of coefficients, except the intercept. Therefore, it substantially reduces the model complexity, and improves the estimation accuracy when the quantiles indeed share similar characteristics (e.g., there are only a few non-zero observations). The option should be used with caution, as its assumption is strong, and it is time consuming.

For the logistic part, L1-penalized logistic regression can be chosen. However, since there are usually adequate data points for the presence-absence model, this option has limited effect in stabilizing estimation.

The final option is to drop the key variables and covariates, and then use the batch ID exclusively in regressions of both parts. We call it simple quantile-quantile matching. In practice, this is essentially the same as matching the empirical quantile functions of each batch to the reference one.

Many factors affect the performance of the standard and alternative fitting strategies, such as the frequencies of taxa, distributional attributes of the read counts (dispersion, heavy tails, or other irregularity), the quality of metadata, etc. Operationally, those options demonstrate different pros and cons for taxa with different frequencies, and the option that is most effective for taxa with a certain range of frequency is data specific. We divide the taxa read count table into sub tables due to frequency, e.g., sub-Tab 1 consisting of taxa with prevalence>90%, sub-Tab 2 consisting of taxa with 80%<prevalence≤90%, etc. For each sub table, we search for the best fitting strategy that produces the least PERMANOVA *R*^2^ explained by batch ID. Concatenating the optimally corrected sub tables, we obtain the overall batch-free microbial profiles. Not that though only local optima (for each sub table) are determined, this procedure is satisfactory considering the extensive time cost by searching for the global optimum (for the overall taxa read count table).

#### Trying over choices of reference batch

ConQuR aligns both the presence-absence likelihood and the distribution of counts given the taxon is present to those of the reference batch. Thus, the quality of the reference batch (both taxa counts and samples metadata) is crucial. Note that a large batch or an abundant batch is not necessarily a high-quality batch. For example, if the reference batch is large, consisting of the most samples, but is extremely sparse, counts of other batches will be drawn to zeros as well; if it is abundant, but mostly taking outlying values, corrected measurements of the other batches might be unstably large.

With each potential reference batch, we conduct the second layer tuning and obtain a corresponding optimally corrected taxa read count table. The corrected table with the least PERMANOVA *R*^2^ is chosen to be the final corrected microbiome data.

### Computation of ConQuR

The computation of ConQuR is intensive as two conditional distributions (original and batch-free) have to be estimated for each sample and each taxon. For a selected fitting strategy, the time depends on sample size *n* and taxa number *J*, and the scale is approximately *O*(*nJ*). Fortunately, the algorithm can be run in parallel because it corrects each taxon separately. In the package, we use two cores to speed it up. For data of similar size as the CARDIA and HIVRC data sets, it will take a PC ~15 min and ~1.75GB memory for correction by standard ConQuR. The complexity increases with tuning. It cost a PC ~2 hours to fine tune the CARDIA or HIVRC data sets over all possible choices. However, in view of the months or years required for data collection, sample processing and bioinformatics, this one-time computation that increases the quality of subsequent analyses should not be a significant concern.

Also, ConQuR is always computationally feasible, regardless of excessive zeros or outlying non-zero observations. This is because it estimates each location of the conditional distribution, separately and non-parametrically. On the other hand, the algorithm for generating negative binomial realizations of ComBat-seq may fail, because taxa that are highly dispersed or with extreme abundances can lead to extraordinarily large estimates of either the mean or dispersion parameters of negative binomial distribution.

## Data availability

The CARDIA data used in this article is available from the CARDIA Study Data Coordinating Center at the University of Alabama at Birmingham. The process for obtaining data through CARDIA is outlined at https://www.cardia.dopm.uab.edu/publications-2/publications-documents. The HIVRC data can be downloaded per instructions in Tuddenham, et al. (2020)^33^.

## Code availability

The R package ConQuR is available at https://github.com/wdl2459/ConQuR in formats appropriate for Macintosh or Windows systems. A vignette demonstrating use of the package is included (can be accessed at https://wdl2459.github.io/ConQuR/ConQuR.Vignette.html).

## Acknowledgements

This work was supported in part by R01 GM129512 and R01 HL155417.

The Coronary Artery Risk Development in Young Adults Study (CARDIA) is supported by contracts HHSN268201800003I, HHSN268201800004I, HHSN268201800005I, HHSN268201800006I, and HHSN268201800007I from the National Heart, Lung, and Blood Institute (NHLBI).

The HIVRC data used in this study are from work that was supported by the HIV Microbiome Re-analysis Consortium. The authors thank Drs. Susan A. Tuddenham and Cynthia L. Sears for their support and review of the manuscript, also thank Dr. Khalil G. Ghanem and all members on the HIV Microbiome Re-analysis Consortium for collecting and processing the HIVRC data set.

## Author contributions

WL developed the method, analyzed the data, and wrote the manuscript. NZ, AMP, WF, AZ, HL, ZL, JC, TR contributed to the conception, methodological developments, and presentation. AL, WLAK, JRW, LJL contributed to the dataset generation and manuscript development with technical input. AAF and KAM contributed to dataset/variable generation, conception, methodological developments, and presentation. MCW conceived the study and critically reviewed the manuscript. All authors read and approved the final manuscript.

## Additional information

Supplementary information is available for this paper in the Supp. file.

## References

1. Lasken, R.S. Genomic sequencing of uncultured microorganisms from single cells. Nature Reviews Microbiology 10, 631–640 (2012).

2. Wooley, J.C., Godzik, A. & Friedberg, I. A primer on metagenomics. PLoS Comput Biol 6, e1000667 (2010).

3. Turnbaugh, P.J. et al. A core gut microbiome in obese and lean twins. nature 457, 480–484 (2009).

4. Qin, J. et al. A metagenome-wide association study of gut microbiota in type 2 diabetes. Nature 490, 55–60 (2012).

5. Mitchell, C.M. et al. Vaginal microbiota and genitourinary menopausal symptoms: a cross sectional analysis. Menopause (New York, NY) 24, 1160 (2017).

6. Langdon, A., Crook, N. & Dantas, G. The effects of antibiotics on the microbiome throughout development and alternative approaches for therapeutic modulation. Genome medicine 8, 1–16 (2016).

7. Claus, S.P., Guillou, H. & Ellero-Simatos, S. The gut microbiota: a major player in the toxicity of environmental pollutants? Npj biofilms and microbiomes 2, 1–11 (2016).

8. Kim, D. et al. Optimizing methods and dodging pitfalls in microbiome research. Microbiome 5, 1–14 (2017).

9. Leek, J.T. et al. Tackling the widespread and critical impact of batch effects in high-throughput data. Nature Reviews Genetics 11, 733–739 (2010).

10. Johnson, W.E., Li, C. & Rabinovic, A. Adjusting batch effects in microarray expression data using empirical Bayes methods. Biostatistics 8, 118–127 (2007).

11. Zhang, Y., Parmigiani, G. & Johnson, W.E. ComBat-Seq: batch effect adjustment for RNA-Seq count data. NAR genomics and bioinformatics 2, lqaa078 (2020).

12. Gibbons, S.M., Duvallet, C. & Alm, E.J. Correcting for batch effects in case-control microbiome studies. PLoS computational biology 14, e1006102 (2018).

13. Dai, Z., Wong, S.H., Yu, J. & Wei, Y. Batch effects correction for microbiome data with Dirichlet-multinomial regression. Bioinformatics 35, 807–814 (2019).

14. Wang, Y. & LêCao, K.-A. Managing batch effects in microbiome data. Briefings in bioinformatics 21, 1954–1970 (2020).

15. Ma, S. et al. Population Structure Discovery in Meta-Analyzed Microbial Communities and Inflammatory Bowel Disease. bioRxiv (2020).

16. Koenker, R. & Bassett Jr, G. Regression quantiles. Econometrica: journal of the Econometric Society, 33–50 (1978).

17. Machado, J.A.F. & Silva, J.S. Quantiles for counts. Journal of the American Statistical Association 100, 1226–1237 (2005).

18. Duan, N., Manning, W.G., Morris, C.N. & Newhouse, J.P. A comparison of alternative models for the demand for medical care. Journal of business & economic statistics 1, 115–126 (1983).

19. Friedman, G.D. et al. CARDIA: study design, recruitment, and some characteristics of the examined subjects. Journal of clinical epidemiology 41, 1105–1116 (1988).

20. Callahan, B.J. et al. DADA2: high-resolution sample inference from Illumina amplicon data. Nature methods 13, 581–583 (2016).

21. Callahan, B. Silva taxonomic training data formatted for DADA2 (Silva version 132). Zenodo (2018).

22. Quinn, T.P. et al. A field guide for the compositional analysis of any-omics data. GigaScience 8, giz107 (2019).

23. Quinn, T.P., Crowley, T.M. & Richardson, M.F. Benchmarking differential expression analysis tools for RNA-Seq: normalization-based vs. log-ratio transformation-based methods. BMC bioinformatics 19, 1–15 (2018).

24. Anderson, M.J. Permutational multivariate analysis of variance (PERMANOVA). Wiley statsref: statistics reference online, 1–15 (2014).

25. Zhao, N. et al. Testing in microbiome-profiling studies with MiRKAT, the microbiome regression-based kernel association test. The American Journal of Human Genetics 96, 797–807 (2015).

26. Huang, J. et al. Six-week exercise training with dietary restriction improves central hemodynamics associated with altered gut microbiota in adolescents with obesity. Frontiers in endocrinology 11(2020).

27. Castelli, W.P. & Anderson, K. A population at risk: prevalence of high cholesterol levels in hypertensive patients in the Framingham Study. The American journal of medicine 80, 23–32 (1986).

28. Ferrier, K.E. et al. Intensive cholesterol reduction lowers blood pressure and large artery stiffness in isolated systolic hypertension. Journal of the American College of Cardiology 39, 1020–1025 (2002).

29. Toya, T. et al. Coronary artery disease is associated with an altered gut microbiome composition. PloS one 15, e0227147 (2020).

30. McInnes, G.T. Hypertension and coronary artery disease: cause and effect. Journal of hypertension. Supplement: official journal of the International Society of Hypertension 13, S49–56 (1995).

31. Pepine, C.J. Systemic hypertension and coronary artery disease. The American journal of cardiology 82, 21–24 (1998).

32. Maifeld, A. et al. Fasting alters the gut microbiome reducing blood pressure and body weight in metabolic syndrome patients. Nature Communications 12, 1–20 (2021).

33. Tuddenham, S.A. et al. The impact of human immunodeficiency virus infection on gut microbiota α-diversity: an individual-level meta-analysis. Clinical Infectious Diseases 70, 615–627 (2020).

34. Daquigan, N., Seekatz, A.M., Greathouse, K.L., Young, V.B. & White, J.R. High-resolution profiling of the gut microbiome reveals the extent of Clostridium difficile burden. NPJ biofilms and microbiomes 3, 1–8 (2017).

35. Vázquez-Castellanos, J.F. et al. Interplay between gut microbiota metabolism and inflammation in HIV infection. The ISME journal 12, 1964–1976 (2018).

36. Kriegeskorte, N., Simmons, W.K., Bellgowan, P.S. & Baker, C.I. Circular analysis in systems neuroscience: the dangers of double dipping. Nature neuroscience 12, 535 (2009).

37. Mullahy, J. Specification and testing of some modified count data models. Journal of econometrics 33, 341–365 (1986).

38. Koenker, R. Econometric Society Monographs: Quantile Regression. New York: Cambridge University (2005).

39. Zou, H. & Yuan, M. Composite quantile regression and the oracle model selection theory. The Annals of Statistics 36, 1108–1126 (2008).

