## Supplementary Materials for "Batch effects removal for microbiome data via conditional quantile regression (ConQuR)"

### Additional details of methods

#### Conditional quantiles of microbial counts $Y_i$

By Model (2), we model a fine sequence of conditional quantiles of $W_i|Y_i > 0$. Due to the one-to-one relationship between quantiles of $W_i|Y_i > 0$ and quantiles of $Y_i|Y_i > 0$[1], together with Model (1), we have the entire conditional quantile function of $Y_i$ piecewise defined,

$$Q_{Y_i}(\tau|\boldsymbol{X}_i) = I\{\tau > 1 - \pi(\boldsymbol{\theta}^L, \boldsymbol{X}_i)\} \cdot \lceil \boldsymbol{X}_i^T \boldsymbol{\theta}^Q \circ \Gamma(\tau; \boldsymbol{X}_i, \boldsymbol{\theta}^L) - 1 \rceil, \qquad (\text{Supp. } 1)$$

where $\pi(\boldsymbol{\theta}^L, \boldsymbol{X}_i) = P(Y_i > 0|\boldsymbol{X}_i)$ and $\Gamma(\tau; \boldsymbol{X}_i, \boldsymbol{\theta}^L)$: $(1 - \pi(\boldsymbol{\theta}^L, \boldsymbol{X}_i), 1) \rightarrow (0,1)$ is a one-to-one mapping from the target quantile level $\tau$ of $Y_i$ to the nominal quantile level $\tau_s$ of $W_i|Y_i > 0$ in Model (2). Specifically, for a $\tau$ below the change point $\pi(\boldsymbol{\theta}^L, \boldsymbol{X}_i)$, the outcome $Y_i$ falls in the "absence range", thus its corresponding quantile is zero. For a $\tau$ beyond the change point, $\Gamma(\tau; \boldsymbol{X}_i, \boldsymbol{\theta}^L)$ tells which quantile of $W_i|Y_i > 0$ determine the value of $Y_i$ at $\tau$. Mathematically,

$$\boldsymbol{\theta}^Q \circ \Gamma(\tau; \boldsymbol{X}_i, \boldsymbol{\theta}^L) = \boldsymbol{\theta}^Q(\tau_s), \tau_s = \Gamma(\tau; \boldsymbol{X}_i, \boldsymbol{\theta}^L) = \frac{\tau - \{1 - \pi(\boldsymbol{\theta}^L, \boldsymbol{X}_i)\}}{\pi(\boldsymbol{\theta}^L, \boldsymbol{X}_i)},$$

and the mapping is derived by

$$\begin{aligned} \tau &= P\{Y_i \leq Q_{Y_i}(\tau|\boldsymbol{X}_i)|\boldsymbol{X}_i\} \\ &= \{1 - \pi(\boldsymbol{\theta}^L, \boldsymbol{X}_i)\} + \pi(\boldsymbol{\theta}^L, \boldsymbol{X}_i) P\{W_i \leq \lceil Q_{W_i}(\tau_s|\boldsymbol{X}_i, Y_i > 0) - 1 \rceil + U|\boldsymbol{X}_i, Y_i > 0\}. \end{aligned}$$

Apart from the characteristics mentioned in the manuscript, another merit of the two-part quantile regression model is that it allows nonlinear associations between quantiles of $Y_i$ and the covariates $\boldsymbol{X}_i$, though the logit of probability being present, and every quantile of the non-zero part are modelled by linear models. To give a concrete example, suppose that for one subject, the true likelihood of the investigated taxon being present in his gut (from Model (1)) is $\pi = 0.8$. Based on the mapping function above, the conditional median abundance of this taxon is determined by the $\tau_s = \frac{0.5-(1-0.8)}{0.8} = 0.38$th quantile of $W_i$. If we change his assignment from placebo to treatment, how would the conditional median of the taxon abundance change accordingly? Suppose the true probability of having the taxon increases to $\pi = 0.9$ after he receives treatment. The conditional median would then be determined by the $\tau_s = \frac{0.5-(1-0.9)}{0.9} = 0.44$th quantile of $W_i$. The difference in conditional median it thus nonlinear, which is a composite effect from the two parts.

**Piecewise estimation strategy**

Parameters of the two-part model (1)(2) can be readily estimated by regressing $I(Y_i > 0)$ on $\boldsymbol{X}_i$ using logistic regression and regressing the non-zero $Y_i$ on the corresponding $\boldsymbol{X}_i$ using linear quantile regression. As the variance of quantile regression estimate is inverse proportional to the local density, that is, $var\{\widehat{\boldsymbol{\theta}}^Q(\tau)\} \to \infty$ as $\tau \to 0$, the quantile estimate is not stable at the change point $1 - \pi(\boldsymbol{\theta}^L, \boldsymbol{X}_i)$, and might blow up to a value with extraordinarily large magnitude. To achieve a reliable estimation of the conditional quantile function $\widehat{Q}$, we use a piecewise strategy:

1. Estimate the probability of presence, $\pi(\widehat{\boldsymbol{\theta}}^L, \boldsymbol{X}_i) = \exp(\boldsymbol{X}_i^T \widehat{\boldsymbol{\theta}}^L)/\{1 + \exp(\boldsymbol{X}_i^T \widehat{\boldsymbol{\theta}}^L)\}$
2. Select a constant $\delta \in (0, \frac{1}{2})$, divide the support of the target quantile levels $(0,1)$ of $Y_i$ into three sub-intervals:

$$A_n = \{\tau: 0 < \tau < 1 - \pi(\widehat{\boldsymbol{\theta}}^L, \boldsymbol{X}_i)\},$$
$$B_n = \{\tau: 1 - \pi(\widehat{\boldsymbol{\theta}}^L, \boldsymbol{X}_i) \le \tau \le 1 - \pi(\widehat{\boldsymbol{\theta}}^L, \boldsymbol{X}_i) + n^{-\delta}\},$$
$$C_n = \{\tau: 1 - \pi(\widehat{\boldsymbol{\theta}}^L, \boldsymbol{X}_i) + n^{-\delta} < \tau < 1\}.$$

3. If $\tau$ is in $B_n$, estimate the quantile coefficients $\widehat{\boldsymbol{\theta}}^Q$ at the nominal quantile level $\Gamma(1 - \pi(\widehat{\boldsymbol{\theta}}^L, \boldsymbol{X}_i) + n^{-\delta}; \boldsymbol{X}_i, \widehat{\boldsymbol{\theta}}^L)$ and perform an interpolation between the quantile

estimate and 1, which is the natural lower bound for microbial read count, at the change point $1-\pi(\widehat{\boldsymbol{\theta}}^L, \boldsymbol{X}_i)$. Pick the interpolated value at $\tau$. If $\tau$ is in $C_n$, estimate the $\widehat{\boldsymbol{\theta}}^Q$ at $\Gamma(\tau; \boldsymbol{X}_i, \widehat{\boldsymbol{\theta}}^L)$ directly. If linear interpolation is used, the estimation of $\hat{Q}$ is shown as Supp. Figure 1, and mathematically is

$$\begin{aligned}
\hat{Q} &= 0 \cdot I(\tau \in A_n) \\
&+ \left[ \{\boldsymbol{X}_i^T \widehat{\boldsymbol{\theta}}^Q \circ \Gamma(1-\pi(\widehat{\boldsymbol{\theta}}^L, \boldsymbol{X}_i) + n^{-\delta}; \boldsymbol{X}_i, \widehat{\boldsymbol{\theta}}^L) - 1\} \cdot \frac{\tau - \{1-\pi(\widehat{\boldsymbol{\theta}}^L, \boldsymbol{X}_i)\}}{n^{-\delta}} \right] \cdot I(\tau \in B_n) \\
&+ \left[ \boldsymbol{X}_i^T \widehat{\boldsymbol{\theta}}^Q \circ \Gamma(\tau; \boldsymbol{X}_i, \widehat{\boldsymbol{\theta}}^L) - 1 \right] \cdot I(\tau \in C_n) \qquad \text{(Supp. 2)}
\end{aligned}$$

Intuitively, the buffer zone $B_n$ avoids the need to estimate the minimal quantiles of the non-zero part, preventing an explosive estimate around the zero-positive change point. The width of $B_n$, $n^{-\delta}$, is designed to converge more slowly than the logistic estimates, so the buffer will work and $\hat{Q}$ is bounded and reliable almost surely. Refer to Ling et al. (2020+)[12] for the consistency of $\hat{Q}$. Other smooth methods can be used instead of the linear interpolation, to further control the estimate around the change point. There is a tradeoff when determining the width of $B_n$. A wider $B_n$ controls the estimates well but the longer interpolation introduces more biases. While if $B_n$ is too narrow, the estimates close to the change point might not be properly controlled. Practically, when the sample size is reasonably large, we use a value approaching $\frac{1}{2}$ as $\delta$, such as 0.499. As a complement to Figure 3b, the matching process on the piecewise estimated conditional quantile functions is depicted in Supp. Figure 2.

However, when the sample size is severely limited, we might opt to omit the buffer zone $B_n$ in estimation. A small sample size leads to a wide $B_n$, then the interpolation in $B_n$ will induce substantial biases. In such extreme cases, reducing the biases is our primary goal, instead of controlling the explosive estimates that might occur.

### References

1. Machado, J.A.F. & Silva, J.S. Quantiles for counts. *Journal of the American Statistical Association* **100**, 1226-1237 (2005).
2. Dillon, S. et al. An altered intestinal mucosal microbiome in HIV-1 infection is associated with mucosal and systemic immune activation and endotoxemia. *Mucosal immunology* **7**, 983-994 (2014).
3. Dinh, D.M. et al. Intestinal microbiota, microbial translocation, and systemic inflammation in chronic HIV infection. *The Journal of infectious diseases* **211**, 19-27 (2015).
4. Lozupone, C.A. et al. Alterations in the gut microbiota associated with HIV-1 infection. *Cell host & microbe* **14**, 329-339 (2013).
5. Monaco, C.L. et al. Altered virome and bacterial microbiome in human immunodeficiency virus-associated acquired immunodeficiency syndrome. *Cell host & microbe* **19**, 311-322 (2016).
6. Noguera-Julian, M. et al. Gut microbiota linked to sexual preference and HIV infection. *EBioMedicine* **5**, 135-146 (2016).
7. Pinto-Cardoso, S. et al. Fecal bacterial communities in treated HIV infected individuals on two antiretroviral regimens. *Scientific reports* **7**, 1-10 (2017).
8. Serrano-Villar, S. et al. The effects of prebiotics on microbial dysbiosis, butyrate production and immunity in HIV-infected subjects. *Mucosal immunology* **10**, 1279-1293 (2017).
9. Vesterbacka, J. et al. (Nature Publishing Group, 2017).
10. Villanueva-Millán, M.J., Pérez-Matute, P., Recio-Fernández, E., Lezana Rosales, J.M. & Oteo, J.A. Differential effects of antiretrovirals on microbial translocation and gut microbiota composition of HIV-infected patients. *Journal of the International AIDS Society* **20**, 21526 (2017).
11. Villar-Garcia, J. et al. Impact of probiotic Saccharomyces boulardii on the gut microbiome composition in HIV-treated patients: a double-blind, randomised, placebo-controlled trial. *PloS one* **12**, e0173802 (2017).
12. Ling, W. et al. Statistical inference in quantile regression for zero-inflated outcomes. *Statistica Sinica*, (2020). In press.

**Figures and tables:**

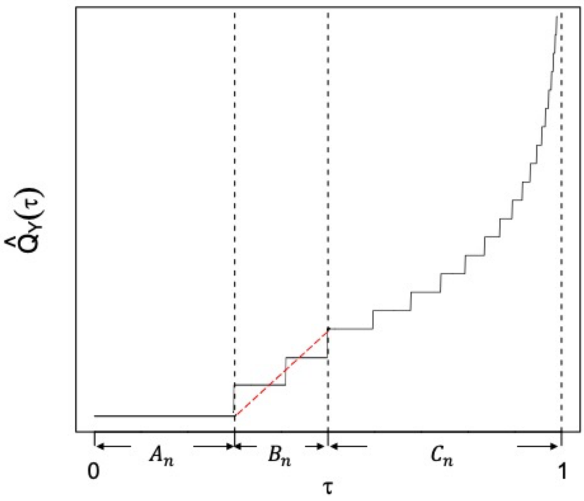


**Supp. Fig. 1 | Piecewise estimation of conditional quantile function.**

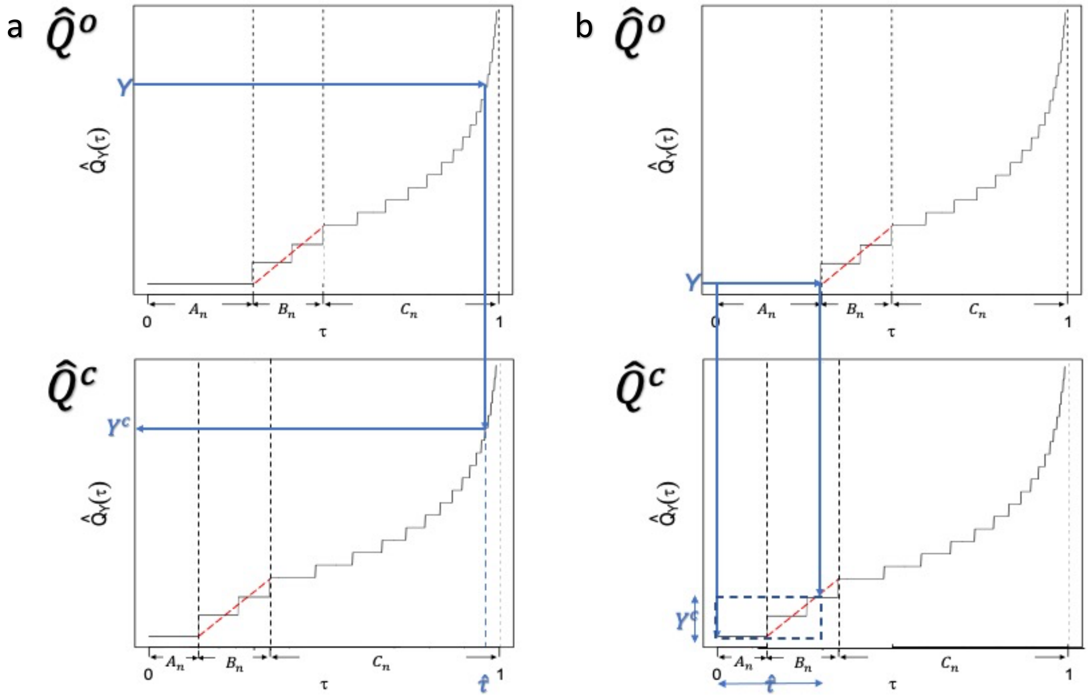

**Supp. Fig. 2 | Correction with conditional quantile functions. a,** Converting a non-zero count to the corrected non-zero count at the same quantile level. **b,** Converting zero to non-zero count when the batch-free distribution is less sparse. Rounded average of all matched quantiles in the batch-free distribution is taken as the adjusted read count.

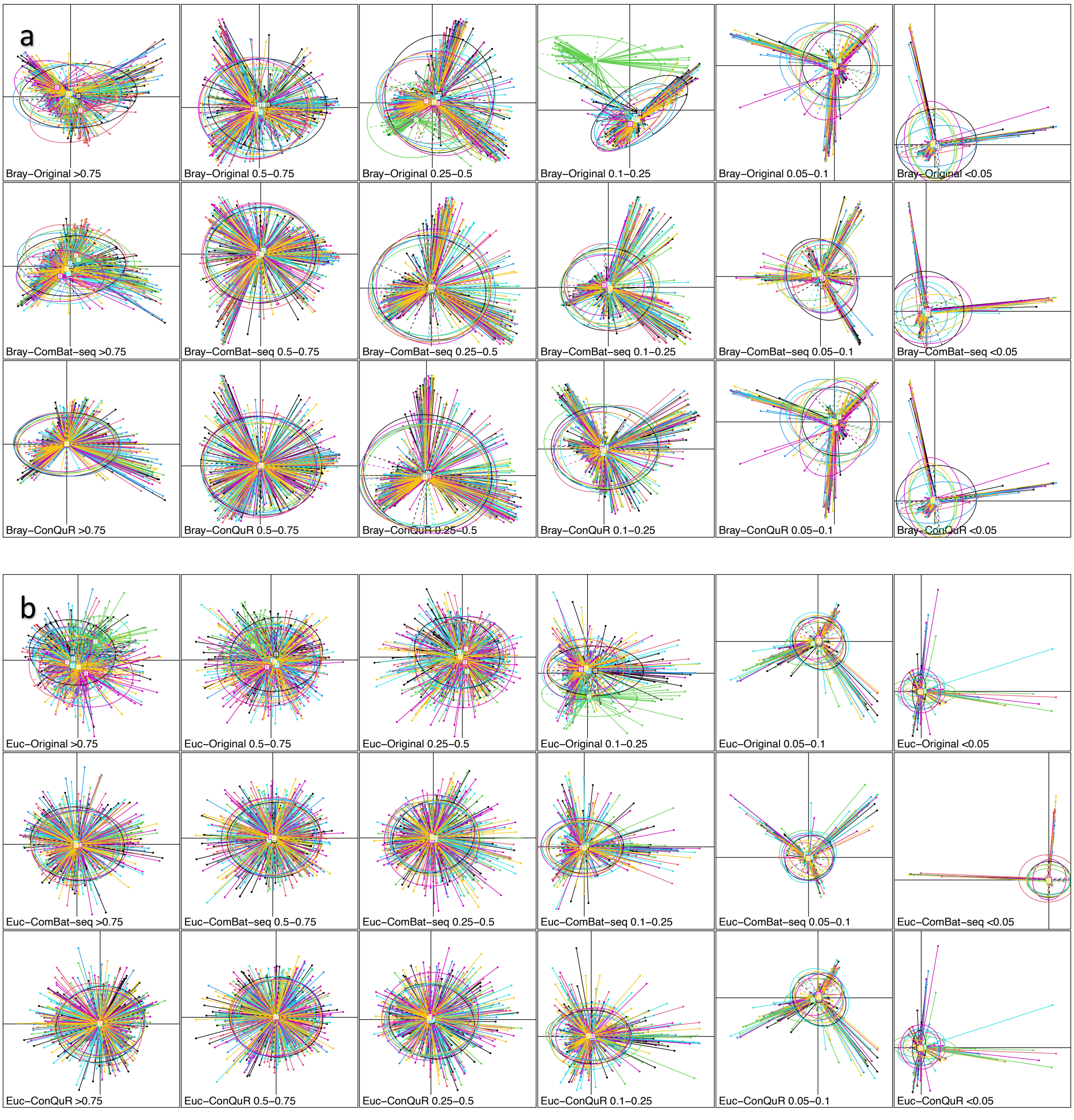

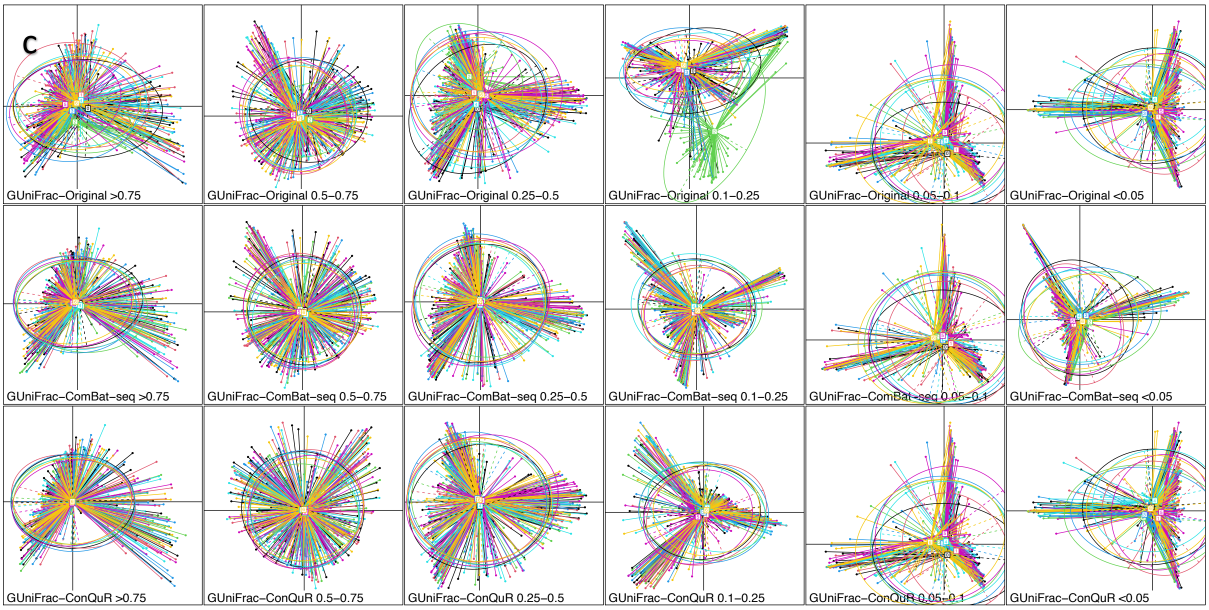


**Supp. Fig. 3 | PCoA plots of original and corrected CARDIA data clustered by batch ID on taxa with different prevalence. a,** by Bray-Curtis distance on original count scale of the raw data, ComBat-seq corrected data, and ConQuR corrected data, on taxa with prevalence >0.75, 0.5-0.75, 0.25-0.5, 0.1-0.25, 0.05-0.1, <0.05. **b,** by Euclidean distance on corresponding CLR transformed relative abundance scale of the data. **c,** by GUniFrac distance.

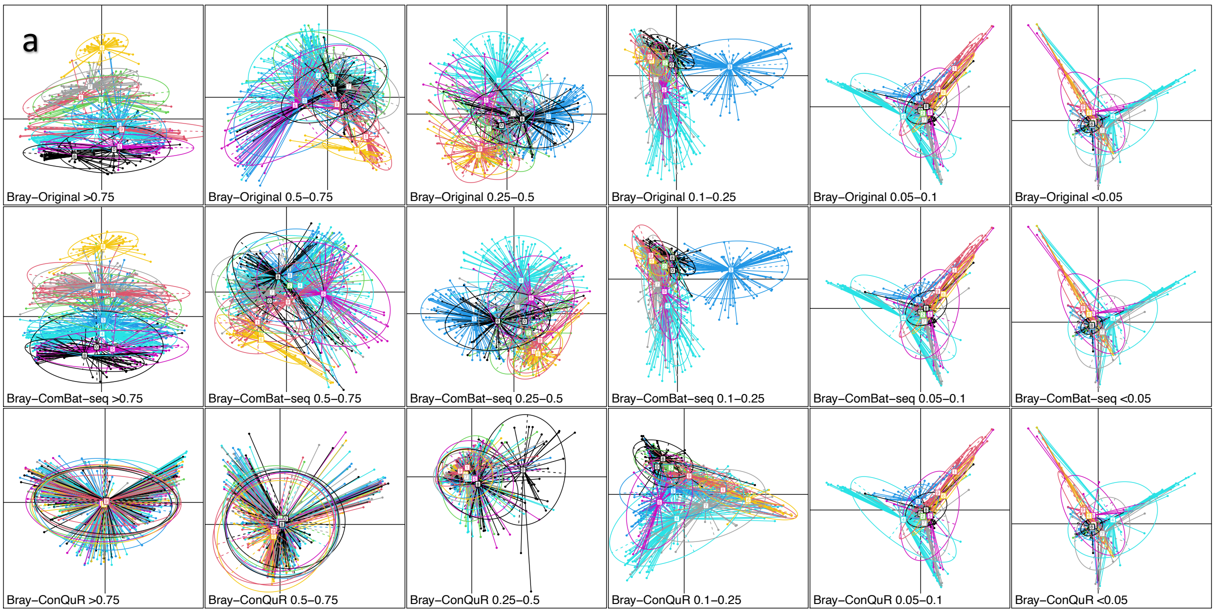

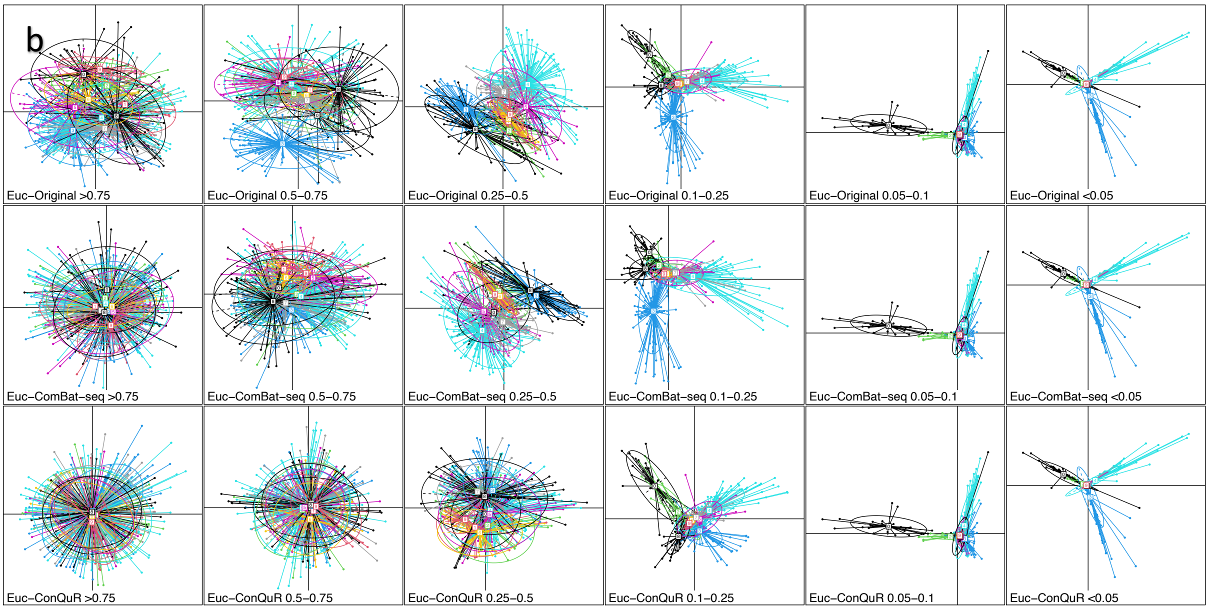


**Supp. Fig. 4 | PCoA plots of original and corrected HIVRC data clustered by study ID on taxa with different prevalence. a,** by Bray-Curtis distance on original count scale of the raw data, ComBat-seq corrected data, and ConQuR corrected data, on taxa with prevalence >0.75, 0.5-0.75, 0.25-0.5, 0.1-0.25, 0.05-0.1, <0.05. **b,** by Euclidean distance on corresponding CLR transformed relative abundance scale of the data.

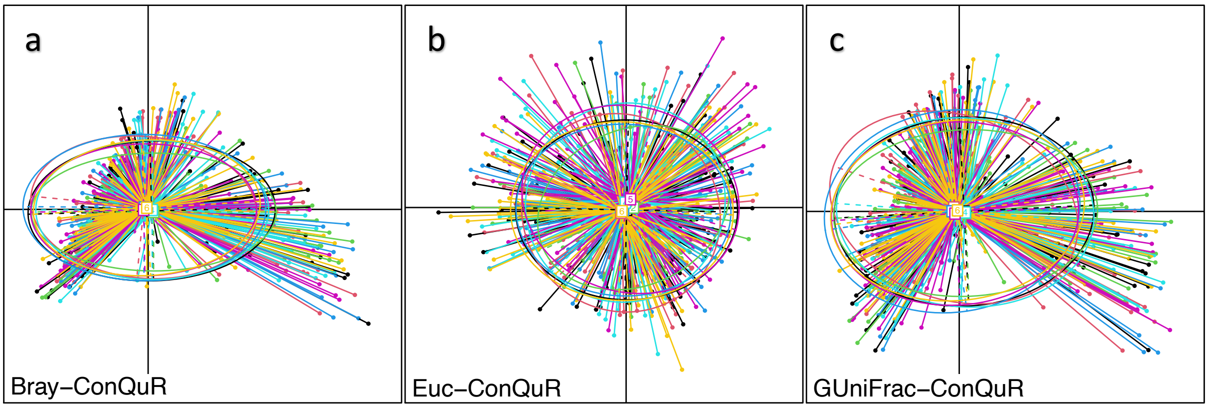


**Supp. Fig. 5 | PCoA plots of corrected CARDIA data by standard ConQuR without fine-tuning, clustered by batch ID. a,** by Bray-Curtis distance on original count scale of the ConQuR corrected data. **b,** by Euclidean distance on corresponding CLR transformed relative abundance scale of the data. **c,** by GUniFrac distance.

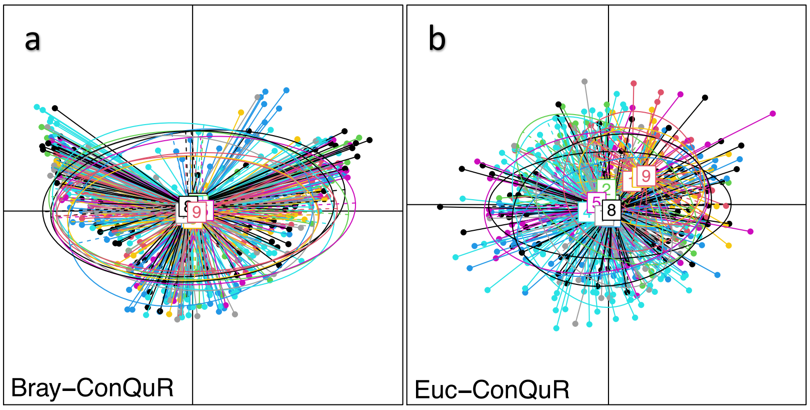


**Supp. Fig. 6 | PCoA plots of corrected HIVRC data by standard ConQuR without fine-tuning, clustered by study ID. a,** by Bray-Curtis distance on original count scale of the ConQuR corrected data. **b,** by Euclidean distance on corresponding CLR transformed relative abundance scale of the data.

**Supp. Tab. 1 | Summary of the 7 runs' microbial profiles and metadata of the CARDIA data set.**

| Batch id | Sample size | Library size | # Genera | SBP<br><br>(mean (SD)) | Gender<br><br>= 1 (%) | Race<br><br>= 1 (%) |
|---|---|---|---|---|---|---|
| 0 | 96 | 67716-139803 | 258 | 118.00 (16.99) | 48 (50.0) | 60 (62.5) |
| 1 | 89 | 53287-117222 | 241 | 117.82 (13.59) | 52 (58.4) | 61 (68.5) |
| 2 | 90 | 61703-112177 | 237 | 121.11 (15.70) | 47 (52.2) | 44 (48.9) |
| 3 | 82 | 69760-124721 | 262 | 119.21 (14.37) | 42 (51.2) | 47 (57.3) |
| 4 | 94 | 84358-193182 | 259 | 119.55 (17.21) | 48 (51.1) | 46 (48.9) |
| 5 | 94 | 90841-224737 | 280 | 121.35 (17.02) | 56 (59.6) | 46 (48.9) |
| 6 | 88 | 71850-253856 | 277 | 117.94 (17.17) | 57 (64.8) | 45 (51.1) |
| | | | Shared: 183 | p-value=0.585 | p-value=0.329 | p-value=0.034 |

**Supp. Tab. 2 | Summary of the 10 sub-studies' microbial profiles and metadata of the HIVRC data set.**

| Study id | Author | Sample size | Library size | # Genera | HIV status = 1 (%) | Age (mean (SD)) | Gender = 1 (%) | BMI (mean (SD)) |
|---|---|---|---|---|---|---|---|---|
| 0 | Dillon[2] | 31 | 20000 | 235 | 18 (58.1) | 35.94 (10.63) | 10 (32.3) | 25.18 (4.44) |
| 1 | Dinh[3] | 36 | 3500 | 157 | 21 (58.3) | 47.66 (7.80) | 8 (22.2) | 25.45 (3.15) |
| 2 | Lozupone[4] | 37 | 5500 | 262 | 24 (64.9) | 36.21 (10.72) | 8 (21.6) | 25.61 (4.84) |
| 3 | Monaco[5] | 110 | 194-10000 | 280 | 73 (66.4) | 39.57 (9.77) | 63 (57.3) | 24.33 (4.48) |
| 4 | Noguera-Julian[6] | 137 | 10000 | 295 | 122 (89.1) | 43.28 (10.12) | 30 (21.9) | 24.15 (3.02) |
| 5 | Pinto-Cardoso[7] | 42 | 15500 | 225 | 33 (78.6) | 40.19 (10.14) | 8 (19.0) | 24.29 (4.09) |
| 6 | Serrano-Villar[8] | 43 | 185-1000 | 143 | 34 (79.1) | 41.79 (10.55) | 7 (16.3) | 24.32 (3.11) |
| 7 | Vesterbacka[9] | 62 | 4000 | 268 | 47 (75.8) | 46.94 (9.61) | 31 (50.0) | 26.45 (4.32) |
| 8 | Villanueva-Millan[10] | 50 | 20000 | 351 | 30 (60.0) | 47.38 (9.49) | 17 (34.0) | 25.73 (5.12) |
| 9 | Villar-Garcia[11] | 24 | 5000 | 148 | 24 (100.0) | 45.92 (9.34) | 3 (12.5) | 23.58 (3.74) |
| | | | | Shared: 65 | p-value<0.001 | p-value<0.001 | p-value<0.001 | p-value=0.004 |
